# Beta band rhythms influence reaction times

**DOI:** 10.1101/2022.11.03.515019

**Authors:** Elie Rassi, Wy Ming Lin, Yi Zhang, Jill Emmerzaal, Saskia Haegens

## Abstract

Despite their involvement in many cognitive functions, beta oscillations are among the least understood brain rhythms. Reports on whether the functional role of beta is primarily inhibitory or excitatory have been contradictory. Our framework attempts to reconcile these findings and proposes that several beta rhythms co-exist at different frequencies. Beta frequency shifts and their potential influence on behavior have thus far received little attention. In this magnetoencephalography experiment, we asked whether changes in beta power or frequency in auditory cortex and motor cortex influence behavior (reaction times) during an auditory sweep discrimination task. We found that in motor cortex, increased beta *power* slowed down responses, while in auditory cortex, increased beta *frequency* slowed down responses. We further characterized beta as transient burst events with distinct spectro-temporal profiles influencing reaction times. Finally, we found that increased motor-to-auditory beta connectivity also slowed down responses. In sum, beta power, frequency, bursting properties, cortical focus, and connectivity profile all influenced behavioral outcomes. Our results imply that the study of beta oscillations requires caution as beta dynamics are multifaceted phenomena, and that several dynamics must be taken into account to reconcile mixed findings in the literature.

## Introduction

Beta rhythms (∼13-30 Hz) are traditionally associated with the sensorimotor system where they are prominent (Pfurtscheller and Lopes da Silva, 1999). Beyond this sensorimotor role, beta has been implicated in a wide range of cognitive phenomena including visual perception (Piantoni et al., 2010; Kloosterman et al., 2015), language processing (Weiss and Mueller, 2012), working memory (Axmacher et al., 2008; Siegel et al., 2009), long-term memory (Hanslmayr et al., 2016), decision-making (Wimmer et al., 2016; Wong et al., 2016), and reward processing (Marco-Pallarés et al., 2015). In non-human primates, beta was shown to reflect top-down attention (Buschman and Miller, 2007), and in rodents beta was linked to working memory (Parnaudeau et al., 2013; Bolkan et al., 2017). However, the functional role of beta is still unclear (Engel and Fries, 2010; Kilavik et al., 2012), as some studies report decreased beta with task engagement, suggesting an inhibitory function, while others report the opposite (Kornblith et al., 2016), suggesting an excitatory function. Similarly, on the neural level, there have been mixed and contradictory findings on the relationship between beta and other neural measures such as firing rate (Rule et al., 2017) and BOLD activity (Hanslmayr et al., 2011).

Current accounts of beta mechanism and function have tried to reconcile these findings (Engel and Fries, 2010; Spitzer and Haegens, 2017). One account states that beta-band activity is related to the maintenance of the current sensorimotor or cognitive state via a top-down mechanism (Engel and Fries, 2010). Our account suggests that beta-band activity is involved in (re)activating latent sensorimotor and cognitive states (Spitzer and Haegens, 2017). We further propose that several beta rhythms co-exist, including functionally inhibitory beta as predominantly observed in sensorimotor regions, and functionally excitatory beta as observed throughout cortex. These different beta rhythms possibly operate at different frequencies (Spitzer and Haegens, 2017). At the neurophysiological level, we posit that while beta events are likely excitatory in nature, there are several biologically plausible ways they could lead to functional inhibition, for example by activating inhibitory neurons or saturating excitatory neurons (Shin et al., 2017; Spitzer and Haegens, 2017).

Beta activity has been characterized and modeled as transient, high-amplitude events or ‘bursts’, which can be detected at the single-trial level (Lundqvist et al., 2016; Sherman et al., 2016). Beta bursts have been observed both focally (Bonaiuto et al., 2021) and as part of long-range communication between brain regions, where beta-band synchrony is assumed to facilitate inter-areal connectivity (Seedat et al., 2020). One property of beta that has received little attention is instantaneous variability in its peak frequency (Cohen, 2014). Here we asked how frequency shifts within the beta band influence behavior.

The influence of beta on behavioral outcomes might depend on several factors such as beta power, frequency, bursting properties, cortical focus, and connectivity profile. In the current experiment we investigated the relationship between single-trial beta activity and behavior, specifically reaction times. Since analyzing neural activity in a pre-stimulus or pre-target interval is a convenient method to uncover the influence of ongoing neural activity on subsequent behavior (Rassi et al., 2019), we made use of magnetoencephalography (MEG) data recorded during an auditory sweep discrimination task. To test how the various characteristics of beta relate to behavior, we analyzed reaction times as a function of pre-target beta differences in power, shifts in frequency, and bursting profiles, within and between motor and auditory cortices.

## Methods

### Participants

We recorded MEG in 35 adult participants, 28 of which we included in our analyses (22 female; mean age = 22.86 years, SD = 2.84; 3 participants excluded due to excessively noisy MEG data and 4 due to near-chance performance on the task). The study was approved by the local ethics committee (CMO Arnhem-Nijmegen). All participants gave informed consent before the experiment and were given monetary compensation for their participation.

### Auditory target discrimination task

The auditory target discrimination task consisted of 5 rhythmic blocks and 5 non-rhythmic blocks (60 trials per block). The order of the blocks was randomized. In the rhythmic blocks, four cue tones were presented, separated by 0.5 s. Following the rhythmic cue, a target tone was presented at 0.5, 1, 1.5 or 2 s (80% of trials) or at .75, 1.25, 1.75 s (20% of trials) after the onset of the last cue tone. In the non-rhythmic blocks, the cue tone was presented continuously for a period of 1.5 s, followed by a target that was presented with a flat probability distribution within a window of 0.5 to 2 s after cue offset. The task for the participant was to determine whether the target tone (a 40-ms chirp) went up or down in pitch. As we had previously shown the experimental cueing manipulation not to produce behaviorally different effects (Wilsch et al., 2020; Lin et al., 2021), here we pooled all trial types (Figure 1a).

**Figure 1.**
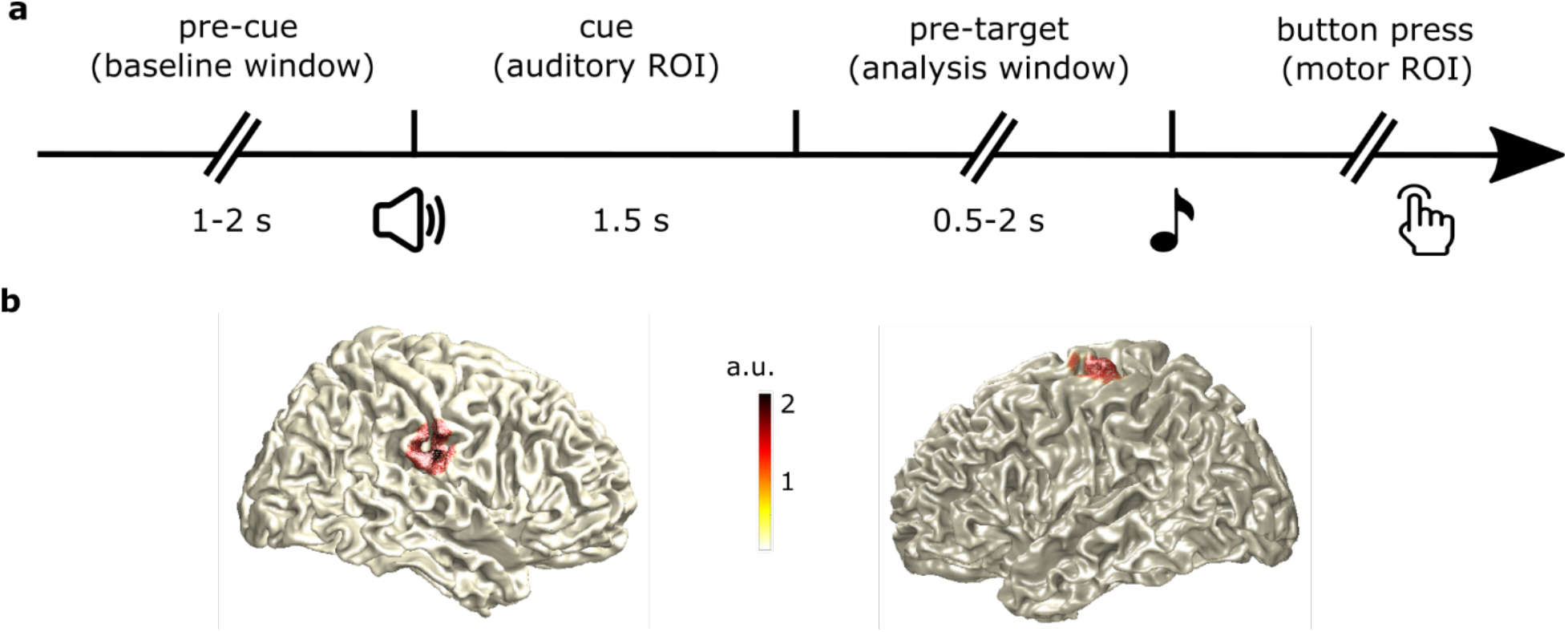
Trial sequence and region of interest definition. a) After a variable baseline delay (1-2 s), an auditory cue lasting 1.5 s played, followed by a variable pre-target delay (0.5-2 s). This pre-target delay was our analysis window. After target onset, participants responded as fast as possible, indicating via button press whether the target tone shifted upward or downward in pitch. b) Regions of interest (ROI) were defined as the source location with maximum evoked activation vs. baseline, based on the evoked response to the auditory cue for the auditory cortex ROI (left panel) and based on the evoked response to the button press for the motor cortex ROI (right). Showing source reconstruction for one representative subject (with a 95%-maximum activity threshold applied for illustrative purposes).

### Stimuli

The cue tones had a pitch frequency of 400 Hz, a sample rate of 44,100 Hz, and a duration of 40 ms (rhythmic blocks) or 1.5 s (non-rhythmic blocks). We used a Hanning taper to remove sharp edges. The target tone consisted of 30 different frequencies randomly drawn from within 500 to 1500 Hz. The target was a frequency-modulated sweep created with the Matlab function *chirp* and was either increasing or decreasing in pitch. The sound had a 10-ms cosine ramp fading in and fading out to avoid onset and offset click perception. The resulting target tone had a sample rate of 44,100 Hz and a duration of 40 ms.

We normalized all sounds (using peak normalization) to the same sound pressure level. We individually adjusted target stimuli to participants’ discrimination threshold, using a custom adaptive-tracking procedure aiming for a discrimination performance between 65 and 85% correct responses. The threshold was the slope of the pitch increase and decrease, measured as the range from lowest to highest frequency (starting point to end). We presented each participant with a pair of sounds (“up” and “down”) consisting of the a priori randomly generated frequencies, modulated depending on their individual threshold; i.e., the 30 base-frequencies were the same for each participant, with the participant’s individual threshold changing the start and end frequencies of the sounds.

### Data acquisition

Whole-head MEG data were recorded at a 1,200-Hz sampling rate with a 275-channel CTF MEG system with axial gradiometers (CTF MEG Systems, VSM MedTech Ltd.) in a magnetically shielded room. To monitor the participants’ head movements online and for offline co-registration of anatomical landmarks, three fiducial coils were placed at the nasion and both ear canals. Anatomical MRI scans for source localization purposes were obtained in a separate session.

### MEG preprocessing

We processed the MEG data offline with the Fieldtrip toolbox (Oostenveld et al., 2011). First, we down-sampled the data to a sampling frequency of 300 Hz. We then applied a notch filter at 50 Hz to remove line noise. Next, we segmented trials into 6-s segments starting 1 s prior to cue onset. We rejected bad channels (∼5%) and bad trials (∼10%) via visual inspection before independent component analysis, which was used to remove components representing eye blinks and heartbeats.

### MEG source reconstruction

We used the obob_ownft toolbox for source reconstruction (https://gitlab.com/obob/obob_ownft). In order to model virtual sensors at the locations of maximum evoked activity in both the auditory and motor sources in the right and left hemispheres respectively, we used a linearly constrained minimum variance (LCMV) beamformer approach (Van Veen et al., 1997). We first constructed volume conduction models of the participants’ brains using a single-shell model of their individual anatomical scans (Nolte, 2003), which we then used to compute leadfields for each gridpoint. Using these leadfields, we computed common spatial filters for each participant using time windows that included a baseline period and the evoked responses.

For the auditory source, we used a time window of 100 ms centered at the peak of the individual auditory evoked response, time-locked to the onset of the auditory cue, and a 100-ms baseline window prior to cue onset. For the motor source, we used an activation time window of 100 ms centered at the peak of the individual motor response, time-locked to the button press, and a 100-ms baseline window prior to the activation window). We then normalized the difference of the sources of the pre and post windows and projected onto their co-registered anatomical scans (Figure 1b). For visualization, we normalized each participant’s brain to Montreal Neurological Institute space. We then identified the location of maximum pre vs. post differences in auditory and motor sources in the right and left hemispheres, respectively. Using the spatial filters for these positions, we then extracted the time series for these two virtual channels.

### Data analysis

We performed all further data analysis using the Fieldtrip toolbox (Oostenveld et al., 2011) and custom Matlab code. We time-locked the source-reconstructed signals from auditory and motor cortices to the onset of the target tones and analyzed a 700-ms pre-target interval. To counteract the 1/f effect in the data, we took the derivative of the time-series data. Note that whether or not we removed the 1/f component had no influence on any of our results. However, beta activity in the resulting flattened spectra was more visually salient, so we used those for visualization.

### Spectral power

To compare pre-target beta power with baseline (i.e., the pre-cue period) activity, and to test the relationship between reaction times and beta power, we extracted 700 ms of pre-cue and pre-target data and computed single-trial Fourier spectra (0-30 Hz) with a fast Fourier approach and a Hanning taper, padded to 2 s for a frequency resolution of 0.5 Hz. We log-transformed the single-trial power data and extracted power in the beta frequency range (13–30 Hz). For the reaction time contrast, we split the pre-target data along the median reaction time, and averaged the power spectra for faster and slower reaction times. We tested for group-level differences in both contrasts with a cluster-based permutation approach (Maris and Oostenveld, 2007), clustering across beta frequencies.

For time-resolved analyses, we performed time-frequency transformation based on multiplication in the frequency domain, using a sliding time window of 250 ms in steps of 20 ms from -750 ms to +250 ms relative to target onset, in steps of 0.5 Hz between 13 and 30 Hz. We then averaged power within this window to obtain a per-region normalization factor and divided each time-frequency point by that factor. Finally, we averaged across the frequency dimension within the beta band and extracted single-trial beta time-courses for the 700-ms pre-target window.

To test the relationship between the beta time-course and reaction time, we z-scored the power values and reaction times, removed those with z-values above 3 and below -3, and used linear regression (Reaction Time = Beta power * slope + intercept), relating each single-trial, pre-target time-point of beta power with the subsequent reaction time on that trial. This provided a time-course of regression slopes per participant. We then generated time-courses of slopes obtained by randomly shuffling the correspondence between power values and reaction times, and tested for group-level differences between the real and shuffled data with a cluster-based permutation approach (Maris and Oostenveld, 2007), clustering across the time dimension (−700 to 0 ms).

### Burst properties

To examine beta burst properties in the source-reconstructed signals, we used the time-frequency representations as described above. We computed the mean and standard deviation of power within a trial for each frequency, and marked the time-frequency points that exceeded two standard deviations above the mean and that lasted at least the duration of one cycle (defined as 1/frequency). We zoomed in on the pre-cue and pre-target delays, and based on temporal and spectral adjacency, we clustered the marked time-frequency points into burst events. We then extracted six parameters of interest from these burst events: For each trial, we counted the number of burst events. Focusing on the event that contained the time-frequency point with the highest power, we extracted the maximum power, the time-point with maximum power, the frequency with maximum power, the frequency range, and the time range. This gave us single-trial estimates of burst properties during the pre-cue and pre-target intervals.

To contrast pre-cue and pre-target burst properties at the group level, we used paired t-tests. To examine the relationship between reaction times and burst properties, we z-scored the burst properties in the pre-target interval and reaction times, and used linear regression analysis to relate them. To test for group-level relationships, we used paired t-tests contrasting the regression slopes against a shuffled distribution.

### Instantaneous frequency

To investigate the time course of the peak beta frequency in the source-reconstructed signals, we analyzed instantaneous frequency as detailed by Cohen (Cohen, 2014). Briefly, we band-passed the single-trial data within the beta frequency range, applied the Hilbert transform, extracted the phase angle time series, took the temporal derivative, and applied ten median filters. This resulted in single-trial time-series of instantaneous frequency during the pre-target interval. To relate those to reaction times, we used the same regression approach detailed above (in Methods, *Spectral power)* to obtain a time-course of regression slopes, and tested them at the group level with a cluster-based permutation approach.

To test the relationship between reaction time and peak frequencies in the power spectra, we averaged power spectra separately for slower and faster trials (based on median split), and detected the peaks of maximum power within the beta range. We then contrasted the peaks at the group level with a paired t-test.

### Connectivity

To estimate the connectivity between auditory and motor cortices, we used the Fourier coefficients that we obtained in the spectral power analysis. As connectivity measures are not resolved on single-trials, we estimated them after splitting the data along the median reaction time. We computed the pairwise phase consistency (Vinck et al., 2010), a bias-free method of rhythmic synchronization. We also computed bi-variate, nonparametric Granger causality (Dhamala et al., 2008a, 2008b), which gave us separate estimates of the connection strengths from motor to auditory cortex and vice versa. We finally contrasted slower vs. faster trials on the group level with a cluster-based permutation approach, clustering across the beta band.

## Results

### Pre-target vs pre-cue power, frequency, and burst properties

First, we examined pre-target beta properties in relation to a baseline (i.e., pre-cue) interval (Figure 2). We found that in both motor and auditory cortex, beta power decreased while beta frequency increased from baseline to pre-target interval.

**Figure 2.**
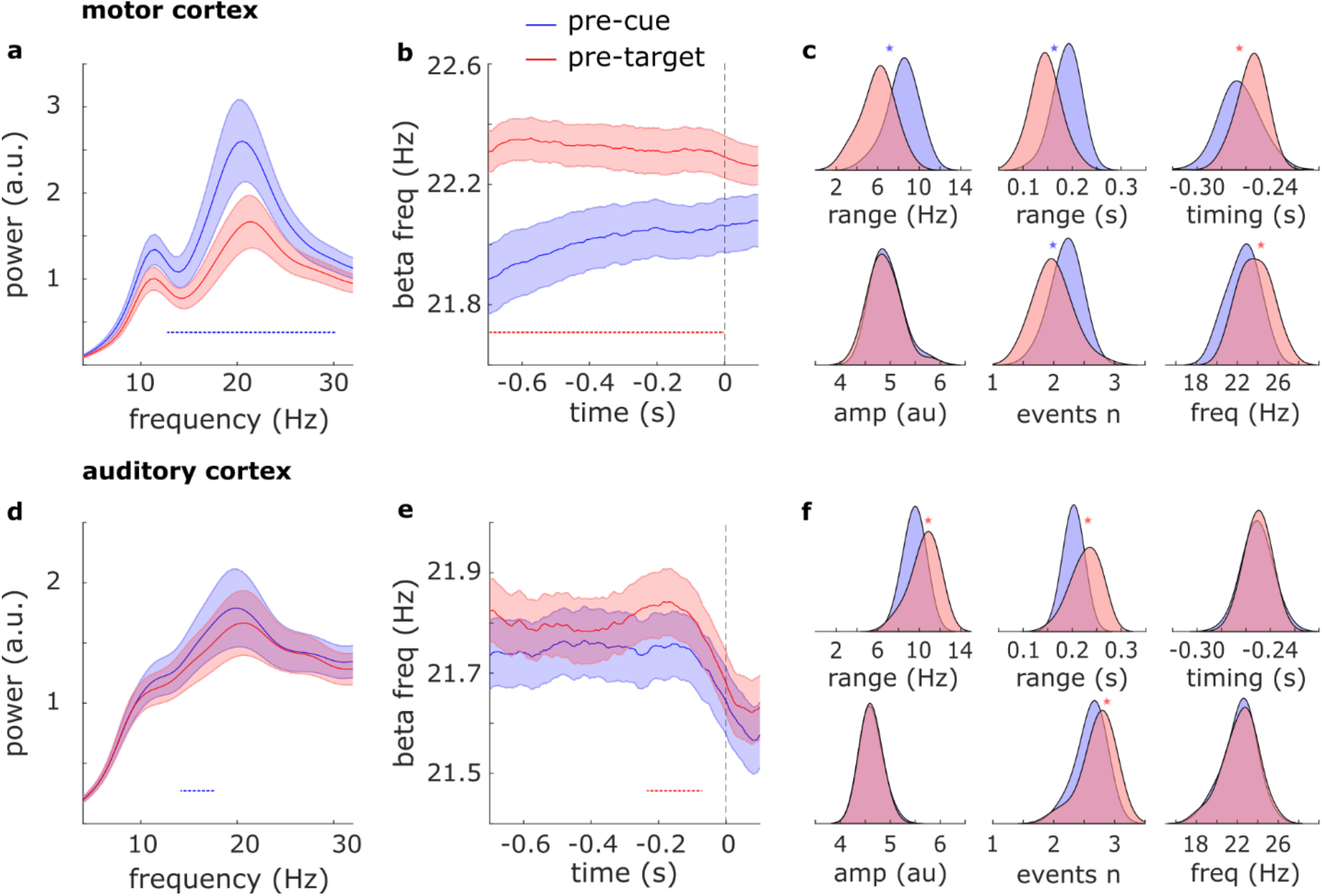
Beta dynamics. a) Power spectra in motor cortex during the pre-target vs. pre-cue delays. b) Instantaneous beta frequency in motor cortex during the pre-target vs. pre-cue delays. c) Beta burst properties in motor cortex during the pre-target vs. pre-cue delays: frequency range, time range, timing relative to target onset, peak amplitude, number of events, and peak frequency. d,e,f) Same as (a,b,c) for auditory cortex. Shaded regions around the line graphs represent the standard error of the mean. Horizontal dotted lines represent significant clusters (p<.05). Spectra in (a) and (d) were detrended by removing 1/f slope. Note that in (b) and (e), the vertical dotted lines corresponding to time-point zero represent the cue onset for the pre-cue time courses (blue) and target onset for the pre-target time courses (red). Asterisks in (c) and (f) represent significant differences between distributions.

In motor cortex, we observed the pre-target power decrease across the whole range of beta frequencies (Figure 2a; cluster-based permutation test across frequencies 13 to 30 Hz; p=1e-5), and the upward shift in beta frequency across the whole interval (Figure 2b; cluster-based permutation test across time -700 to 0 ms; p=2e-4). Consistently, there were fewer bursts (t(27)=-5.8, p=3.4e-6), with narrower time spans (t(27)=-9.8, p=2e-10) and narrower frequency spans (t(27)=-12.3, p=1.4e-12), and the peak burst frequency was also increased (t(27)=5.8, p=3.7e-6) during the pre-target delay as compared to baseline (Figure 2c). In addition, bursts in motor cortex happened closer in time to target onset than they did to cue onset (t(28)=2.8, p=.009). There were no differences in the maximum power of the bursts.

In auditory cortex, we observed the pre-target beta power decrease primarily in the 14.5 to 17 Hz range (Figure 2d; p=.019), and the upward shift in beta frequency primarily from 210 to 75 ms prior to target onset (Figure 2e; p=.039). However, there were more bursts (t(27)=5.3, p=1.3e-5) with wider time spans (t(27)=5.7, p=5e-6) and wider frequency spans (t(27)=6.0, p=1.9e-6) during the pre-target delay as compared to baseline (Figure 2f). Note that while a power decrease seems inconsistent with more bursts, the power decrease occurred at frequencies lower than the frequencies at which bursts occurred (i.e., 14.5 to 17 Hz vs. ∼22 Hz), and lower than peak beta frequency (which was at 20 Hz based on peak detection, or 22 Hz based on instantaneous frequency analysis), making this decrease in power hard to interpret. There were no differences in the maximum power of the bursts, their peak frequency, or their timing relative to stimulus onset.

Next, we tested whether pre-target beta properties related to reaction times using two complementary approaches: a median-split approach to relate beta measures to slow vs. fast reaction times, and a regression approach to relate single-trial beta measures to reaction times. The two approaches yielded the same results: in motor cortex, slower reaction times were related with higher beta power, while in auditory cortex, slower reaction times were related with higher beta frequency.

### Spectral power

In motor cortex (Figure 3 a,b,c), slower reaction times were preceded by higher beta power. Splitting the power spectra across the median reaction time revealed the effect was driven by differences in the 20 to 26 Hz frequency range (Figure 3a; cluster-based permutation test across frequencies: p=1e-5). This effect was present throughout the whole pre-target interval when looking at the time-resolved power envelopes (Figure 3b; cluster-based permutation test across time: p=6e-4). A time-resolved single-trial regression approach confirmed this effect as well (Figure 3c; slope = 0.030; cluster-corrected p=4e-4). To further characterize this difference, we zoomed in on the beta bursting profile and found that slower reaction times were preceded by more bursts (slope = 0.040; p = .0017) with wider time spans (slope = 0.037; p = .0277) and wider frequency spans (slope = 0.039; p = .0143).

**Figure 3.**
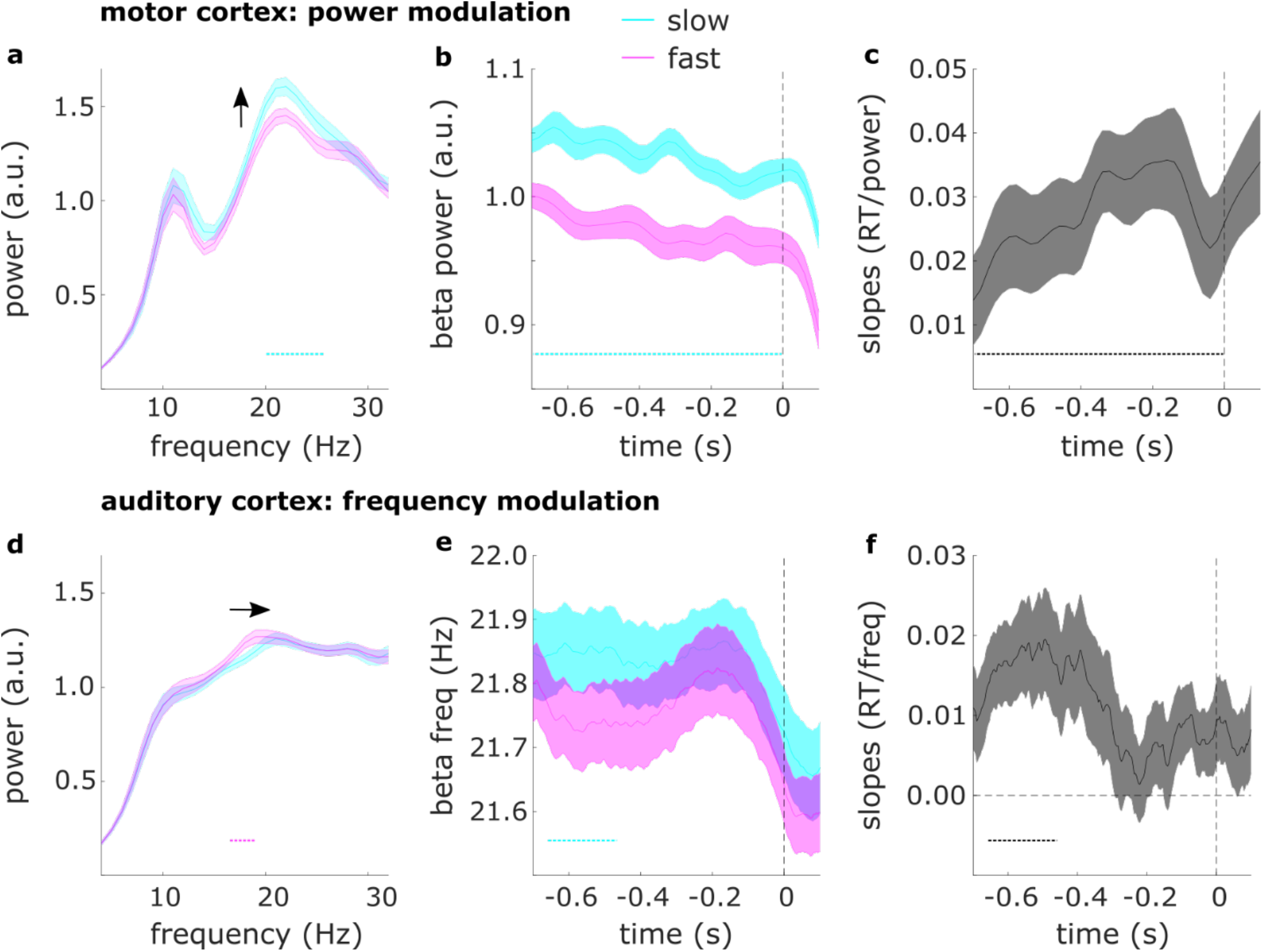
Relation between beta dynamics and reaction times. a) Power spectra in motor cortex for trials with slow vs. fast reaction times. b) Time-resolved beta power in motor cortex for trials with slow vs. fast reaction times. c) Regression slopes for the relationship between reaction times and time-resolved beta power in auditory cortex. d) Same as (a) for auditory cortex. e) Instantaneous beta frequency in auditory cortex for trials with slow vs. fast reaction times. f) Regression slopes for the relationship between reaction times and instantaneous frequency in auditory cortex. Shaded regions around the line graphs represent the standard error of the mean. Horizontal dotted lines represent significant clusters (p<.05). Vertical dashed lines represent time-point zero (target onset). Spectra in (a) and (d) were detrended by removing 1/f slope.

In auditory cortex, when splitting the data along the median reaction time and contrasting the power spectra, we found an effect opposite to that observed in motor cortex, such that faster reaction times were preceded by higher beta power (Figure 3d; cluster-corrected p=.016), an effect driven by differences in the 17 to 19 Hz frequency range. However, pre-target beta power was not related to reaction times when using the time-resolved regression approach (slope = -0.008, no clusters). Given the discrepancy in results between the two approaches, we further investigated the observed difference in auditory cortex as a possible shift in peak frequency.

### Beta frequency

In auditory cortex (Figure 3 d,e,f), slower reaction times were preceded by a higher peak beta frequency when splitting the power spectra across the median reaction time and detecting participants’ individual peak beta frequencies (t(24)=2.4, p=.0255). This effect was most pronounced around 660 to 450 ms prior to target onset when looking at instantaneous frequency (Figure 3e; cluster-based permutation test across time: p= .006). A time-resolved single-trial regression approach confirmed the same result (Figure 3f; slope = 0.011; cluster-corrected p=.032). When zooming in on the peak burst frequencies, we found the same relationship again (slope = 0.029; t(27) = 2.45; p = .021). In motor cortex, beta frequency was not related to reaction times (slope = -0.0052; no significant clusters).

### Connectivity

We then asked whether auditory-motor beta connectivity was related to reaction times (using the median-split approach). Slower reaction times were preceded by increased beta connectivity between auditory and motor cortices, as quantified with pairwise phase consistency (p = .027). The difference was most prominent at frequencies from 19 to 20 Hz (Figure 4a). We then used Granger causality to check the directionality of this effect. There were no differences in auditory-to-motor beta connectivity (Figure 4b; no significant clusters), but slower reaction times were preceded by increased motor-to-auditory beta connectivity (p = .047), most prominently at frequencies from 20 to 21 Hz (Figure 4c).

**Figure 4.**
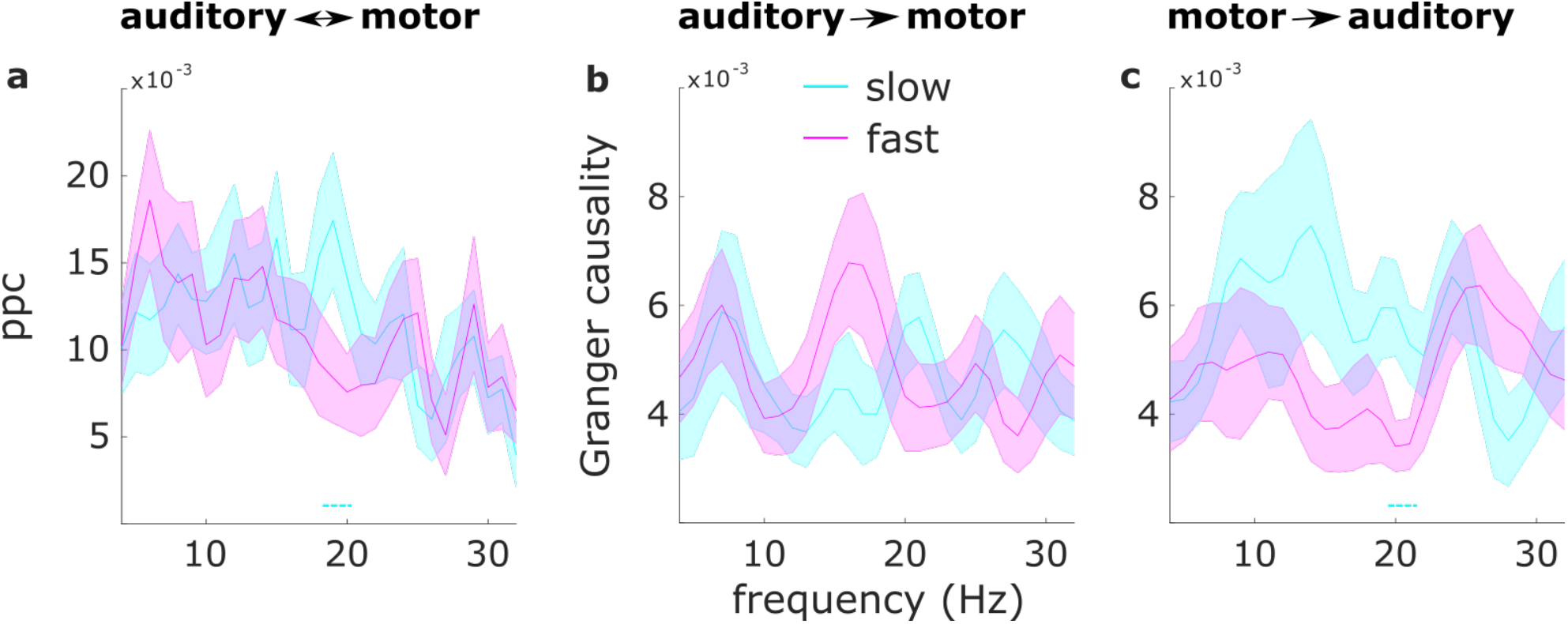
Auditory-motor cortex connectivity. a) Pairwise phase consistency between auditory and motor cortices for slow vs. fast reaction times. b) Granger causality from auditory to motor cortex for slow vs. fast reaction times. c) Granger causality from motor to auditory cortex for slow vs. fast reaction times. Shaded regions around the line graphs represent the standard error of the mean. Dotted lines represent significant clusters (p<.05).

## Discussion

In an auditory target discrimination task, we sought to uncover the relationship between reaction times and various characteristics of the beta rhythm. We found that slower (as compared to faster) reaction times were preceded by increased beta power in motor cortex, increased beta frequency in auditory cortex, and increased motor-to-auditory connectivity in the beta range. The results were robust across our analysis approaches. We used a regression approach to relate single-trial reaction times to beta measures, as well as a median-split approach to relate slower vs. faster trials to changes in beta measures, with both approaches yielding the same pattern of results. We further analyzed beta activity separately in a time-resolved manner, a frequency-resolved manner, and by characterizing its burst profile, with all approaches yielding the same pattern of results.

An analysis of the burst properties of the increased beta activity prior to slower responses revealed that this increased activity likely reflected an increased number of pre-target bursts with wider time and frequency ranges. These results are in line with an account of beta which is bursting in nature and inhibitory in function (Shin et al., 2017). However, our power results in auditory cortex are at odds with this inhibitory function. At the single-trial level, pre-target auditory beta power was not robustly related to reaction times. Based on this set of results, we conclude that the beta rhythm potentially serves different functions in different cortical locations. The beta rhythms in motor and auditory cortex differed not only in their impact on behavior, but also in their peak frequencies. Although a crude distinction between ‘higher’ and ‘lower’ beta across cortical locations has been previously made (e.g., Kopell et al., 2011), attempts to assign them different functional roles have had mixed success (Spitzer and Haegens, 2017).

One aspect of beta rhythms (and cortical rhythms in general) that has so far received little attention is the non-stationarity of their frequency across time. Models of beta function account for the potential of different beta rhythms occurring at different frequencies, assuming different cortical locations or different generators within a location. But it is so far under-appreciated that a single rhythm can shift in frequency over time (Cohen, 2014). Frequency shifts according to task demands have been observed in human EEG/MEG data for the alpha rhythm (Haegens et al., 2014; Samaha and Postle, 2015; Wutz et al., 2018), and in non-human primate LFP data for the beta rhythm (Kilavik et al., 2012). It has also been reported that slower alpha rhythms correlate with slower responses across subjects (Surwillo, 1961), but to our knowledge the relationship between beta frequency and reaction times has not yet been investigated. We here report the opposite relationship for the beta rhythm, such that faster (auditory) beta correlated with slower reaction times within subjects.

Beyond local beta dynamics, beta has also been shown to be involved in long-range communication between cortical sites (Seedat et al., 2020). Here we found increased beta connectivity between motor and auditory cortex, specifically in the direction of motor to auditory cortex, prior to slower (vs. faster) responses. It is unlikely that this effect was confounded by the power difference in motor cortex as we used a phase-based connectivity measure, and in addition, there were no robust power differences in auditory cortex. This finding is in line with the notion of covert active sensing, where the motor system actively coordinates sensory systems (Schroeder et al., 2010). Our effects can also be interpreted in light of the frequency-matching notion (Lowet et al., 2017). That is, on trials with slow responses, the difference in peak frequencies between auditory and motor cortex is reduced, resulting in stronger synchronization. This interpretation is supported by our observation of stronger auditory-motor coupling on slow trials (Figure 2).

Finally, our results imply that the analysis of beta oscillations requires caution as beta dynamics are multifaceted phenomena. For example, it is possible that observed power modulations are better explained as frequency shifts (as is the case for our results). It is also possible that the beta rhythm serves different functions (i.e., inhibitory or excitatory) depending on the cortical region where it is found or depending on whether it is local or inter-areal. Future investigations could focus on inter-areal variability in beta peak frequency, for example in intracranial human electrophysiological recordings.

## Author contributions

ER conceived the study, analyzed the data, and wrote the manuscript. WML contributed to data analysis and writing. YZ contributed analysis code. JE collected the data. SH conceived the study and wrote the manuscript.

## Acknowledgements

We would like to thank Anna Wilsch and Iske Bakker-Marshall for assistance during data collection, and Julio Rodriguez-Larios for valuable comments on the manuscript. This work was supported by an FWF Erwin-Schroedinger fellowship J480 to ER, and by NWO grants Veni 451-14-027 and 016.Vidi.185.137 to SH.

## Data and code availability

All data and code supporting the findings of this study will be made publicly available upon acceptance of the manuscript.

